# *Fusobacterium necrophorum* and *Fusobacterium varium* are commensal members of the bovine reproductive microbiota and may colonize calf prenatally

**DOI:** 10.1101/2024.10.15.618546

**Authors:** Justine Kilama, Carl R Dahlen, Mina Abbasi, Xiaorong Shi, T. G. Nagaraja, Matthew S. Crouse, Robert A. Cushman, Alexandria P. Snider, Kacie L. McCarthy, Joel S. Caton, Samat Amat

## Abstract

*Fusobacterium necrophorum* is an important pathogen associated with several infectious diseases in cattle. However, recent sequencing-based studies have indicated that *F. necrophorum* is positively associated with pregnancy in beef cows and that *Fusobacterium* is the most abundant genus in the bull seminal microbiota, suggesting the potential role of *Fusobacterium* in reproductive health and fertility. Here, we performed a comprehensive screening to 1) determine whether *Fusobacterium necrophorum* (subspecies *necrophorum* [FNN] and *funduliforme* [FNF]), and *Fusobacterium varium* (FV) are part of the commensal members of the reproductive microbiota in cattle; 2) to explore whether these *Fusobacterium* spp. are colonized in calf prenatally. For this, we screened 11 different sample types including bovine and ram semen, bovine vaginal and uterine swabs, and bull fecal samples, as well as samples from 180- and 260-days old calf fetuses and their respective dams using both quantitative PCR (514 samples) and targeted culturing (499 samples). By qPCR, all the targeted *Fusobacterium* spp. were detected across all sample types, with FNF being the highly prevalent in the bull semen (66.7%) and maternal ruminal fluids (87.1%), which was confirmed by culturing. All the targeted *Fusobacterium* were identified in vaginal and uterine (3.1%-9.4%) as well as placental caruncles, and fetal fluids, ruminal and meconium samples (2.7% - 26.3%) by qPCR and were not isolated by culture method. Overall, our results suggest that *F. necrophorum* is a commensal member of healthy male reproductive microbiota, and that FNF, FNN and FV are present in bovine vagino-uterine microbiota, and calf intestine prenatally.

## INTRODUCTION

Traditionally, the microbes in reproductive tracts of mammals have been predominantly regarded as pathogens, with the semen and uterus long believed to be sterile, and any microbial presence considered detrimental [1–5]. However, the advent of high-throughput sequencing techniques has altered this perspective, revealing that the reproductive tract hosts diverse microbial communities that are important in reproductive health and fertility [2, 6]. Moreover, the core microbial community of a healthy bovine reproductive tract and its state of balance with the host described as *eubiosis* is yet to be clearly defined [7]. This is partly because the reproductive microbiota is potentially shaped by a range of animal-specific factors, including stage of the estrous cycle, pregnancy status, timing relative to parturition, parity, breed, and genetics [8, 9], as well as herd management practices such as nutrition, calving assistance, and housing conditions [9]. For instance, while *Fusobacterium* species (spp.) is mostly recognized as a major pathogen responsible for a wide spectrum of infections in both animals and humans [10], its potential involvement in bovine reproductive health appears to be more complex.

*Fusobacterium necrophorum* is a Gram-negative, rod-shaped, non-spore-forming, non-motile, pleomorphic, aerotolerant anaerobe that can colonize the gastrointestinal, respiratory, and reproductive tracts of both humans and animals [11–13]. *F. necrophorum* is classified into two subspecies (subsp.) namely: subsp. *necrophorum* (FNN) and subsp. *funduliforme* (FNF) [14]. These subspecies exhibit distinct genetical [13], morphological, biochemical, and virulence characteristics [15], with FNN being notably more virulent than the FNF owing to its ability to produce more leukotoxin [16]. Recently, *F. varium,* which seemingly share similar characteristics with *F. necrophorum,* in terms of habitat and biochemical niche such as indole production and the utilization of lactate and lysine [17–19], has been identified as the predominant *Fusobacterium* spp. in the bovine ruminal microbial ecosystem [20–22].

In addition to liver abscesses, necrotic laryngitis and foot rot, *F. necrophorum* has been associated with mastitis, and abortion in cattle [10, 12, 23]. *Fusobacterium* spp. have also been implicated in the development of endometritis, especially in transition dairy cows experiencing metabolic stress and immunological impairments [7, 24, 25]. Thus, *F. necrophorum* is considered to be one of the most important bacterial pathogens implicated in many infectious diseases in cattle [26, 27] and can contribute to the increased antimicrobial usage and significant economic losses to both dairy and beef cattle industries. However, recent emerging body of evidence derived from culture-independent sequencing-based studies indicate that *Fusobacteirum* spp., including *F. necrophorum,* might be a commensal member of the microbial communities associated with healthy male [28] and female reproductive tracts [29]. Our previous 16S rRNA gene amplicon sequencing-based study revealed that the phylum Fusobacteriota accounts for 6.3% and 0.6% of the vaginal and uterine microbiota of healthy beef heifers, respectively [29]. The genus *Fusobacterium* was found in vagina and uterus of these healthy heifers with noticeably high abundance [29]. In another study, we characterized the uterine microbiota of beef cows at the time of artificial insemination (AI) using 16S rRNA gene sequencing and identified 11 amplicon sequencing variants (ASVs) including *F. necrophorum* which were more abundant in the uterine microbiota of beef cows that became pregnant than that of those cows that did not become pregnant [30]. This observation suggests the potential positive association of *F. necrophorum* with cattle fertility.

A number of 16S rRNA gene amplicon sequencing based studies, including our own, revealed that Fusobacteriota is one of the most predominant bacterial phyla comprising the seminal microbiota of healthy beef bulls [28, 31–34]. We recently identified *Fusobacterium* as the most dominant genus (26% relative abundance) in the seminal microbiota of healthy beef bulls [28], and observed that its relative abundance was reduced in the bull semen during breeding, suggesting the potential transfer from the bull semen to the female reproductive tract. This evidence coupled with the increased appreciation of the role of urogenital microbiota in male and female reproductive health and fertility [35], as well as possible hitch-hiking of the microbes with the semen to the female reproductive tract [4, 36] together prompted us to investigate whether *F. necrophorum* subspecies and *F.varium* are commensal members of the bovine reproductive microbiota. The primary objective of this study was to screen and identify FNN, FNF, and FV in male and female reproductive tracts of bovine and ovine using qPCR and targeted culturing methods. The secondary objective was to examine the presence of *Fusobacterium* spp. in calf fetuses and their respective dams at late gestation, to explore their potential prenatal colonization in cattle. Unlike earlier studies that predominantly utilized 16S rRNA gene amplicon sequencing which is often limited to characterizing microbiota mainly at genus level, this study employed both culture-based and qPCR-based techniques that specifically focused on *Fusobacterium* spp. This approach enabled both quantification, identification and characterization of *Fusobacterium* at the species and subspecies levels which provide better understanding of the prevalence of *Fusobacterium* spp. in bovine reproductive tract. By including large number of samples derived from many different animal trials, this study establishes a foundational framework for future investigations into potential involvement of *Fusobacterium* spp. in cattle fertility and the fetal immune programming.

## MATERIAL AND METHODS

All animal handling and experimental procedures in this study were approved by the relevant institutional committees for animal research. The protocols involving bulls were approved by the Institutional Animal Care and Use Committee (IACUC) for the U.S. Meat Animal Research Center (USMARC IACUC; experiment number 147.2). The rest of the protocols were authorized by the IACUC in the North Dakota State University (NDSU), including IACUC protocol #A20059 for rams, protocol #A21061 for cows and heifers sampled at the time of artificial insemination. Procedures involving late gestational calves, and their dams were approved under NDSU protocol IACUC20210043 for 180-days old fetuses and IACUC ID # A21049 for 260-days old fetuses.

### Study designs, animal husbandry and dietary treatments

To approach this research topic with a broad scope of sample types and animal species, we utilized samples from five different studies involving beef bulls, rams, beef heifers and cows, as well as 180- and 260-day-old beef calf fetuses and their respective dams as illustrated in Fig.1. While the diets and animal husbandry and management practices were varied between different animal studies, sample collections and processing for microbial analysis from all the animal studies were conducted following the same procedures by the same personnel.

**Figure 1:**
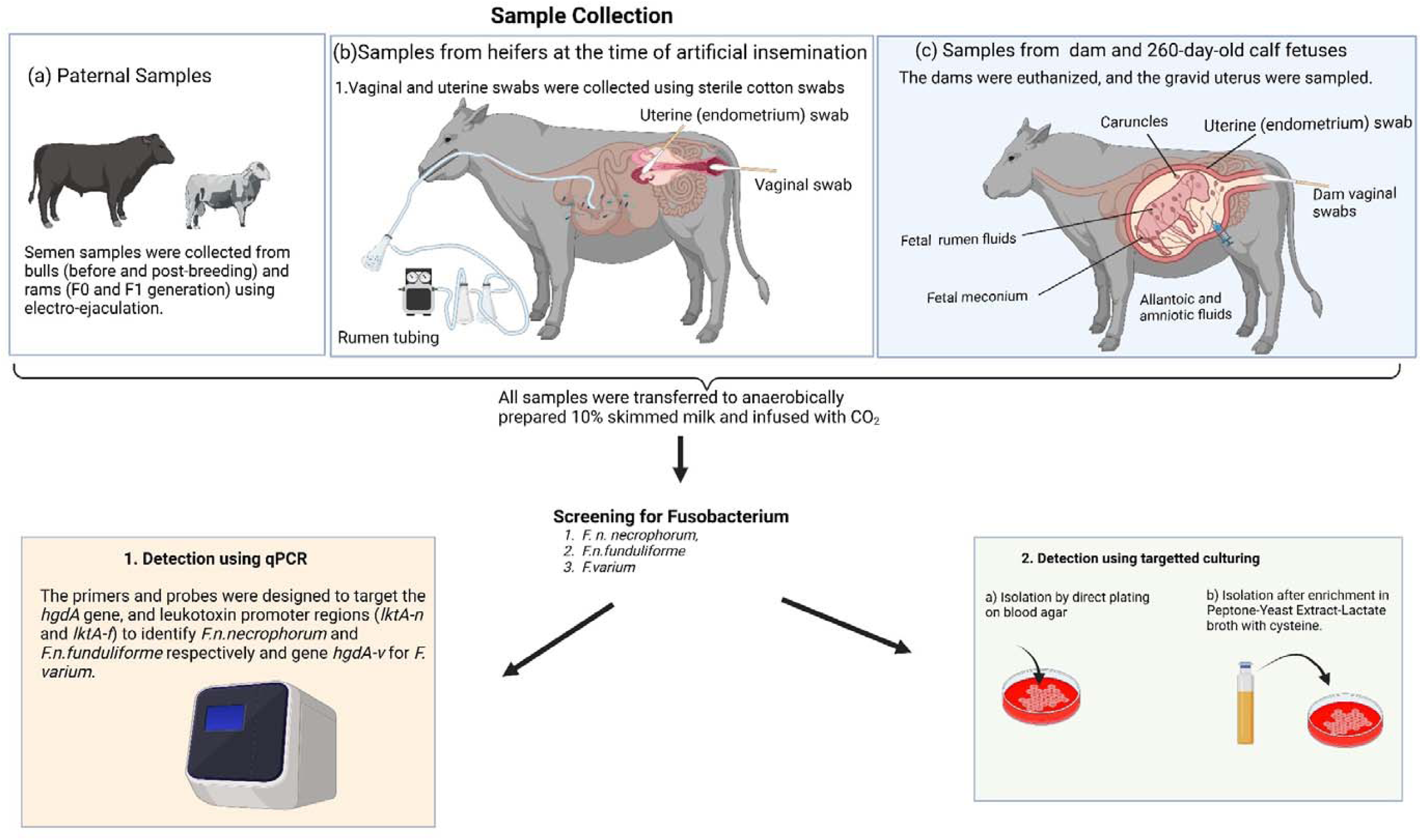
Illustration of experimental design, sample collection procedures and screening techniques for *F.necrophorum* and *F. varium*

The first study involved beef bulls with cohorts conducted in each of the three consecutive years, from 2022 to 2024. The detailed husbandry practices used for managing the bulls are described in our previous publication [28]. In brief, 40 MARC II bulls, aged 9-10 months old, were subjected to either moderate (1.13 kg/day) or high (1.80 kg/day) average daily weight gain feeding regimens over 112 days, with biweekly adjustments to feed delivery to ensure targeted BW gain was achieved. The diet consisted of 25% alfalfa hay, 5% corn silage, 66% corn, and 4% vitamin/mineral pellets, administered via Calan gates.

The second study involved rams, with a primary goal of evaluating the impacts of sire plane of nutrition on offspring outcomes. The study design with the husbandry practices and dietary management are described previously [37]. Briefly, 24 Rambouillet rams, aged 1.5 to 4 years, with an initial average body weight of 82.9 ± 2.6 kg, were acclimated for 16 days at the NDSU Animal Nutrition and Physiology Center, where they were housed individually in temperature-controlled pens with simulated natural daylight. Over an 84-day feeding period, the rams were assigned to one of three nutritional planes: Positive, aiming for a 12% gain in body weight; Maintenance, designed to maintain their current body weight; and Negative, targeting a 12% reduction in body weight. The F1 rams were fed a common diet for 7 months when semen samples were collected. Semen samples used in this study were collected from the F0 generation at different sampling days (days 0, 28,56, 84) and from their male offspring at one sampling time.

The third study involved a herd of Angus-crossbred cows (27 open, 31 pregnant) from which both uterine and vaginal swabs were collected at time of AI as well as heifers (26 open, 33 pregnant) from which only vaginal swabs were collected 2 days before AI to characterize the reproductive microbiota and its relation to pregnancy outcomes. A detailed description of the study is described in our previous publication [30].

The fourth study consisted of 20 heifers which were managed on either a high forage diet (75% forage, 25% concentrate) or a high concentrate diet (75% concentrate, 25% forage) at the NDSU Beef Cattle Research Complex. The high forage diet comprised 54% alfalfa hay, 42% corn silage, and 4% vitamin/mineral pellet (dry matter basis), while the high concentrate diet included 10% alfalfa hay, 30% corn silage, 56% corn, and 4% vitamin/mineral pellet (dry matter basis). The heifers were bred with artificial insemination using male-sexed semen and maintained on their respective dietary treatments until 180 days of gestation, at which time they were slaughtered and tissue from dams and fetuses were collected (unpublished data).

The fifth study included a herd of 32 pregnant heifers, which were fed either control or restricted diets, with some receiving one-carbon metabolites (OCM) supplementation, including methionine, choline, folate, and vitamin B12 as described previously [38]. The heifers were inseminated with female-sexed semen and maintained on OCM dietary treatments until 260 days of gestation, at which time they were slaughtered and tissue from dams and fetuses were collected (unpublished data).

### Sample collection

#### a) Bull semen and fecal sample collection (first study)

Bull semen (n = 36) was collected during pre-breeding soundness exams after day 112 of being managed on pasture and post-breeding, following a 28-day breeding season with 250 heifers [39]. Briefly, semen samples were collected through electroejaculation, using a collection handle lined with a plastic sheath (Pulsator IV; Lane Manufacturing Inc.; Denver, CO) as outlined by [28]. A 200 µL aliquot of the bull semen were transferred into cryotubes with 1 mL of 10% skim milk saturated with carbon dioxide gas and immediately placed on dry ice. Fecal samples were collected from the bulls from the rectum (∼150 grams) using plastic glove (without additional lubrication) and placed into sterile Whirl-Pak bags. Approximately 5 mg of feces were then transferred into 1 mL of 10% skim milk (Difco^TM^ Skim milk, Sparks, MD, USA) that was saturated with carbon dioxide gas. Both semen and fecal samples were frozen immediately and transported to the lab on dry ice for storage at −80°C until further analysis.

#### b) Ram semen sample collection (second study)

Ram semen was collected on days 0, 28, 56, and 84 using electroejaculation, with samples collected into a handle lined with a plastic sheath [37]. A 50 µL portion of the semen was added to 1 mL of brain heart infusion (BHI) broth with 20% glycerol for culturing and additional 200 µL of the semen was stored in a 1.5 mL tube for qPCR assay following genomic DNA extraction.

#### c) Sampling maternal ruminal fluid, uterine and vaginal swabs from live animals (third study)

Maternal ruminal fluid was collected from cows using a pump-assisted rumen tube [29], aliquoted into cryotube containing 10% skim milk (DifcoTM Skim milk) and immediately placed on dry ice before transportation to the laboratory for processing. The vaginal and uterine swabs collections were detailed in our previous publication [30]. Briefly, the vulva was thoroughly cleaned with a sterile paper towel soaked in 70% ethanol. A sterile cotton-tipped applicator (15 cm, Puritan, Guilford, ME, USA) was then inserted into the midpoint of the vaginal cavity and swirled four times to ensure adequate contact with the vaginal wall and was withdrawn carefully to avoid contamination. The swab was immediately placed into a cryotube with 10% skim milk (Difco^TM^ Skim milk), stored on ice, and transported to the laboratory. Immediately after collecting the vaginal swabs, uterine swabs were obtained using a 71 cm double-guarded culture swab (Reproduction Provisions L.L.C., Walworth, WI, USA), which was carefully guided through the cervix into the uterine body with the aid of rectal palpation. The swab tip was extended through the inner plastic sheath, gentle pressure was applied to the uterine body by pinching the swab and rotating it three times. The swab was then retracted into its sheaths and removed from the cow. The tip of the swab was cut off, placed into a 2-mL tube with 10% skim milk (Difco^TM^ Skim milk), kept on ice, and transported to the laboratory for processing.

#### d) Collection of dam and fetus-associated samples (fourth and fifth study)

Samples collected from the dam consisted of ruminal fluids, vaginal swabs, and uterine swabs, while fetus-associated samples included allantoic and amniotic fluids, placental caruncle tissues, ruminal fluids, and meconium. Immediately post-euthanasia via captive bolt, maternal ruminal fluid samples were collected after evisceration of gastrointestinal tract. About 50 mL of ruminal fluid were drawn into a sterile 50 mL syringe (Medtronic, Minneapolis, MN) using a sterile 22-gauge needle. The cranial vagina was swabbed as described above, and uterine swabs were obtained after exposing the gravid uterus from the dam during fetus harvesting. All samples were stored in 10% skim milk infused with carbon dioxide gas prior to immediate cryopreservation for subsequent DNA extraction and culturing.

As illustrated in Fig. 1, various fetus-associated samples, including allantoic and amniotic fluids, caruncles, ruminal fluids, and meconium were collected under aseptic conditions. Allantoic and amniotic fluid samples were obtained as detailed in our previous publications [40, 41]. Briefly, immediately after exposing the gravid uterus, up to 50 mL of allantoic and amniotic fluid were aspirated using a sterile 22-gauge needle into a sterile 50 mL syringe. Fetal rumen fluid was collected using a procedure similar to that used for maternal ruminal fluid collection as described above. A 500 µL aliquot of each amniotic, allantoic, and rumen fluid sample was transferred into 1 mL of 10% skim milk infused with carbon dioxide gas and immediately placed on dry ice. Soon after opening through the uterine walls, and separation from the fetal cotyledon, about 5.0 mg of maternal placental caruncle tissue were dissected [40], immersed in 1 mL of 10% skim milk infused with carbon dioxide gas, and placed on dry ice. Fetal meconium samples were collected into sterile Whirl-Pak bags, and a sterile spatula was used to transfer about 5.0 mg sample into 1 mL of 10% skim milk infused with carbon dioxide gas, followed by immediate freezing on dry ice. To monitor potential contamination, sterile swabs were used to sample surgical trays, tables, instruments, room air, and tap water from the abattoir floor. The number of each type of sample collected and utilised for either qPCR or targeted culturing are summarised in Table 1. Samples were preserved on dry ice and transported to the Department of Diagnostic Medicine and Pathobiology at Kansas State University (Manhattan, KS, USA) for qPCR analysis and targeted culture method.

**Table 1:** Sample size distribution of different types of samples screened for *F. necrophorum* and *F. varium* using qPCR and culturing methods.

| Sample type | Number of samples screened |  |
| --- | --- | --- |
|  | qPCR | Culturing |
| Bull semen | 96 | 116 |
| Bull feces | 24 | 24 |
| Bovine maternal ruminal fluid | 31 | 31 |
| Bovine vaginal swab | 55 | 68 |
| Bovine uterine swabs | 64 | 79 |
| Bovine caruncles | 38 | 38 |
| Bovine allantoic fluids | 34 | 34 |
| Bovine amniotic fluids | 37 | 37 |
| Fetal ruminal fluid | 31 | 31 |
| Fetal meconium | 31 | 31 |
| Ram semen | 100 | 10 |
| **Total number of samples** | **541** | **499** |

### Genomic DNA extraction

DNA extraction was conducted using the GeneClean Turbo Kit (MP Biomedicals, Solon, OH) as previously described by [21]. Briefly, 1 mL of each homogenized samples were boiled for 10 minutes and centrifuged at 9,300 × g for 5 minutes. The supernatant (100 μL) was mixed with 500μL of GeneClean Turbo Salt Solution and transferred to a cartridge, then centrifuged at 14,000 × *g* for 5 seconds. After washing and additional centrifugation, DNA was eluted with 30μL of GeneClean Turbo Elution Solution following a 5-minute incubation at room temperature and a final one-minute centrifugation. The eluted DNA was stored at −20°C until qPCR analysis.

### Quantitative polymerase chain reaction (qPCR)

A total of 541 genomic DNA samples, extracted from the 11 different sample types, before and after enrichment (described below), were analysed using qPCR (Table 1) to detect and quantify the two subspecies of *Fusobacterium* and *F. varium* [21]. The *hgdA* gene, whih encodes 2-hydroxyglutaryl dehydratase, was used for identification of *F. necrophorum* (*hgdA-n*) and *F. varium* (*hgdA-v*), and the promoter region (*lktA-n* and *lktA-f*) of the leukotoxin operon (*lktBAC*) was used to differentiate between the two subspecies of *F. necrophorum* [42]. The targeted species or subspecies in different sample types were considered present when qPCR assay was positive either pre or post enrichment. Conversely, it was considered not detected if the qPCR assay yielded negative both prior to and after enrichment.

### Targeted Culturing

#### Enrichment media

The media used to enrich samples were pre-reduced, anaerobically sterilized peptone yeast extract medium (PY) with 100 mM lactate (PY-La) or lysine (PY-Ly) as the primary carbon source and supplemented with josamycin (3 μg/mL), vancomycin 4 μg/mL, and norfloxacin 1 μg/mL; PY-La JVN or PY-Ly JVN; [21, 22] prior to isolation.

#### Isolation of Fusobacterium *spp*

A total of 499 samples (Table 1) were subjected to targeted culturing both by direct plating without enrichment and plating after enrichment in PY-La-JVN or PY-Ly-JVN. A detailed protocol for isolation of *Fusobacterium* spp. has been previously described [22]. In short, sample homogenates were spot inoculated onto blood agar (Remel Inc., Lenexa, KS), PY-La JVN agar, and PY-Ly JVN agar using sterile cotton swabs. Subsequently, an inoculating loop was used to streak from the inoculation spot to facilitate the isolation of single bacterial colonies. These inoculated plates were incubated in an anaerobic glove box at 37°C for 48 hours. Presumptive colonies, based on morphology shown in Fig. 3A [22] were selected and transferred onto blood agar plates and incubated anaerobically at 37°C for 48 hours. Confirmation of the species identification was performed by a qPCR assay targeting the *hgdA* gene. For enrichment, 1 mL of sample homogenate was inoculated into PY-La JVN and PY-Ly JVN broths, incubated at 37°C for 24 hours, and then streaked onto blood agar for isolation. The sample was considered positive for *Fusobacterium* spp. or subspecies, if presumptive colonies exhibiting the morphological characteristics of the *Fusobacterium* species or subspecies were positive by qPCR.

### Statistical Analysis

The results of qPCR assay and targeted culturing performed both before and after enrichment were individually combined to calculate the total prevalence of FV, FNN, and FNF in each sample type, The cumulative prevalence data derived from these two detection methods were then compiled and displayed in figures created with Prism software (Prism 9.5.0, GraphPad).

## RESULTS

In the present study, we investigated the prevalence of FV, FNN, and FNF in the male and female reproductive tracts of cattle and sheep utilizing diverse sample types including vaginal and uterine samples from beef heifers and cows as well semen samples from beef bulls and rams (Fig. 1). Additionally, we evaluated the presence of *Fusobacterium* spp. in late-gestational calf fetuses and their dams to explore their prenatal colonization.

### Detection of *Fusobacterium* spp. by qPCR

Of the 541 samples subjected to qPCR before and after enrichment, FNF was the most prevalent *Fusobacterium* subspecies in majority of sample types analyzed (Fig. 2). The concentrations ranged from 1 × 10³ to 1.5 × 10LJ CFU per mL, with an average concentration of 7 × 10³ CFU per mL. The highest prevalence of FNF was in maternal ruminal fluids, with a prevalence of 87.1% followed by bull semen at 66.7%. FNF was also detected in 21.1% of bovine caruncles, and 16.1% of bovine fetal ruminal fluids, 5.9% of allantoic fluids, 5.5% of bovine maternal vaginal swabs, and 3.1% of uterine swabs. However, FNF was not detected in bull fecal samples, amniotic fluids, fetal meconium, and ram semen samples. Enrichment revealed the presence of FNF in bovine uterine swabs and fetal ruminal fluid.

**Figure 2:**
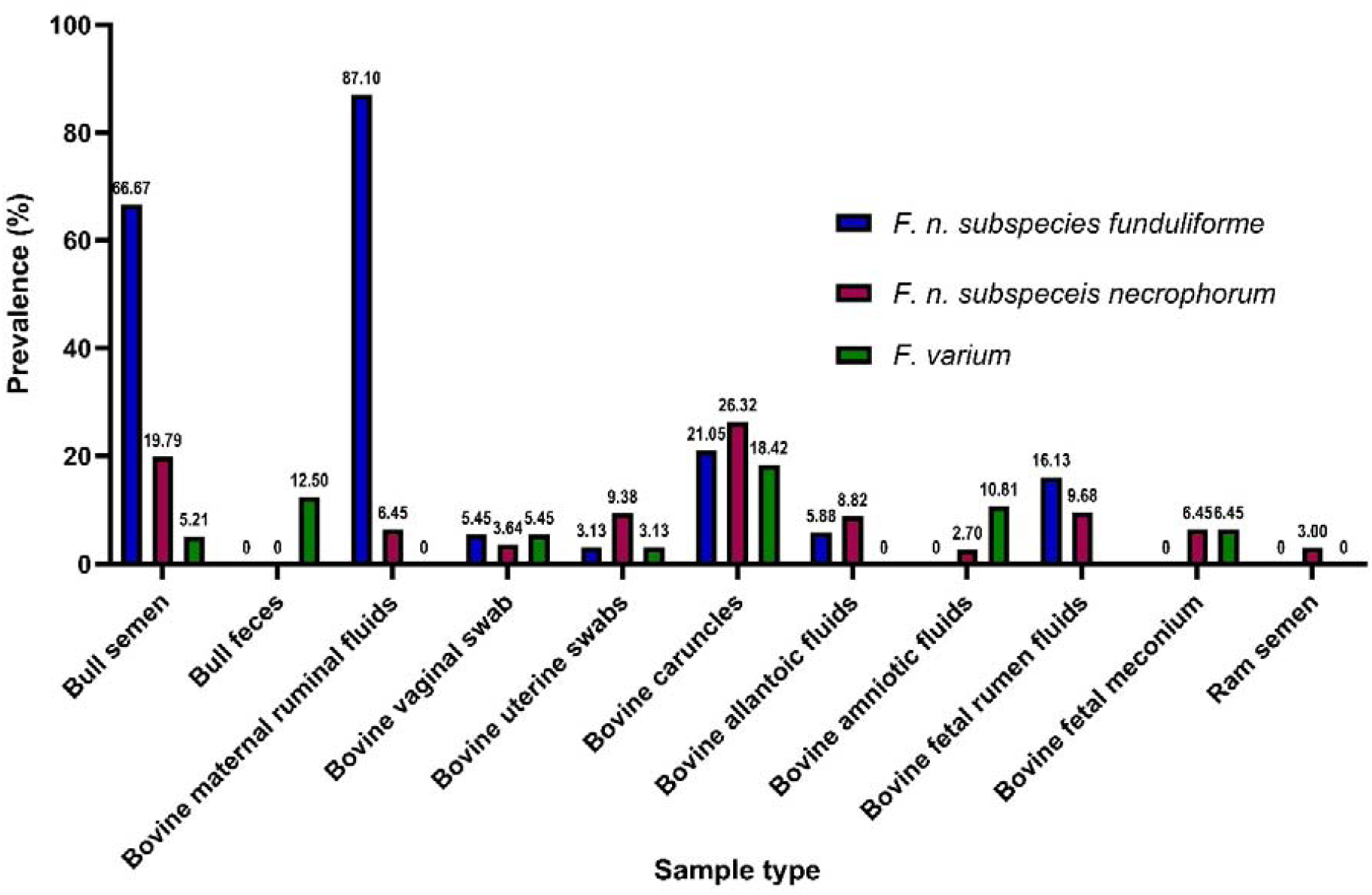
Total prevalence of targeted *Fusobacterium* spp. in different sample types (n=541) as screened using qPCR.

**Table 2:** Frequency of detection of targeted *Fusobacterium* spp. in different sample types using qPCR both before and after enrichment.

| Sample type | N | *F. n. subspecies funduliforme* |  |  | *F. n. subspecies necrophorum* |  |  | *F. varium* |  |  |
| --- | --- | --- | --- | --- | --- | --- | --- | --- | --- | --- |
|  |  | Before enrichment <sup>a</sup> | After enrichment <sup>a</sup> | Total prevalence (%) | Before enrichment <sup>a</sup> | After enrichment <sup>a</sup> | Total prevalence (%) | Before enrichment <sup>a</sup> | After enrichment <sup>a</sup> | Total prevalence (%) |
| Bull semen | 96 | 25 (26.0%) | 39 (54.9%) | 64 (66.7%) | 7 (7.3%) | 12 (13.5%) | 19 (19.8%) | ND | 5 (5.2%) | 5 (5.2%) |
| Bull feces | 24 | ND | ND | ND | ND | ND | ND | ND | 3 (12.5%) | 3 (12.5%) |
| Bovine maternal ruminal fluids | 31 | 1 (3.2%) | 26 (86.7%) | 27 (87.1%) | ND | 2 (6.5%) | 2 (6.5%) | ND | ND | ND |
| Bovine vaginal swab | 55 | 1 (1.8%) | 2 (3.7%) | 3 (5.5%) | ND | 2 (3.6%) | 2 (3.6%) | 1 (1.8%) | 2 (3.7%) | 3 (5.5%) |
| Bovine uterine swabs | 64 | ND | 2 (3.1%) | 2 (3.1%) | 1 (1.6%) | 5 (7.9%) | 6 (9.4%) | ND | 2 (3.1%) | 2 (3.1%) |
| Bovine caruncles | 38 | ND | 8 (21.1%) | 8 (21.1%) | 5 (13.2%) | 5 (15.2%) | 10 (26.3%) | ND | 7 (18.4%) | 7 (18.4%) |
| Bovine allantoic fluids | 34 | ND | 2 (5.9%) | 2 (5.9%) | ND | 3 (8.8%) | 3 (8.8%) | ND | ND | ND |
| Bovine amniotic fluids | 37 | ND | ND | ND | ND | 1 (2.7%) | 1 (2.7%) | 4 (10.8%) | ND | 4 (10.8%) |
| Fetal ruminal fluid | 31 | ND | 5 (16.1%) | 5 (16.1%) | ND | 3 (9.7%) | 3 (9.7%) | ND | ND | ND |
| Fetal meconium | 31 | ND | ND | ND | ND | 2 (6.4%) | 2 (6.4%) | ND | 2 (6.4%) | 2 (6.4%) |
| Ram semen | 100 | ND | ND | ND | 3 (3.0%) | ND | 3 (3.0%) | ND | ND | ND |
| **Total** | **541** | **27 (5.0%)** | **84 (15.5%)** |  | **16 (3.0%)** | **35 (6.5%)** |  | **5 (0.9%)** | **21 (3.9%)** |  |

<sup>a</sup> Cultured for 24 hours in a peptone yeast extract medium, utilizing either lactate (100 mM) or lysine (100 mM) as the main energy source, and supplemented with josamycin (3 µg/mL), vancomycin (4 µg/mL), and norfloxacin (1 µg/mL), ND=Not detected, N=Number of samples.

In contrast, FNN had a lower prevalence compared to FNF although it was found across all sample types examined, except in bull feces. The highest prevalence of FNN was in bovine caruncles at 26.3%, followed by 19% in the bull semen. It was also present in 9.7% of fetal ruminal fluids, 9.4% of uterine swabs, 8.8% of allantoic fluids, and 6.5% of both maternal ruminal fluids and fetal meconium. Additionally, FNN was detected in 3.6% of maternal vaginal swabs, 3.0% of ram semen and 2.7% of amniotic fluids.

Lastly, FV was the least prevalent of the *Fusobacterium* species across all the samples screened. It was detected most frequently in bovine caruncles (18.4%), followed by bull faeces (12.5%) and amniotic fluids (10.8%). FV was also found in 6.5% of fetal meconium, 5.5% of bovine maternal vaginal swabs, 5.2% of bull semen, and 3.1% of uterine swabs (Fig.2). However, it was undetectable in bovine allantoic fluids, as well as in maternal and fetal ruminal fluids, and ram semen, even after enrichment.

### Isolation of *Fusobacterium* spp

A total of 499 samples were subjected to culturing in order to isolate FNN, FNF and FV by direct plating or plating after an enrichment step. Consistent with the qPCR results, FNF was the most frequently isolated subspecies. The greatest prevalence of FNF was in the maternal ruminal fluid, where it was isolated from 67.7% of samples, followed by the bull semen (62.9%) (Fig. 3B). However, FNF was less frequently isolated from other sample types, with only a single isolate recovered from 68 bovine vaginal swabs, and none from uterine swabs, caruncles, allantoic fluids, amniotic fluids, fetal rumen fluid, fetal meconium, and ram semen (Fig. 3B). In bull semen, the number of isolates increased by 23 following enrichments, while maternal ruminal fluid and vaginal swabs yielded an additional 14 and 1 isolates, respectively (Table 3).

**Figure 3:**
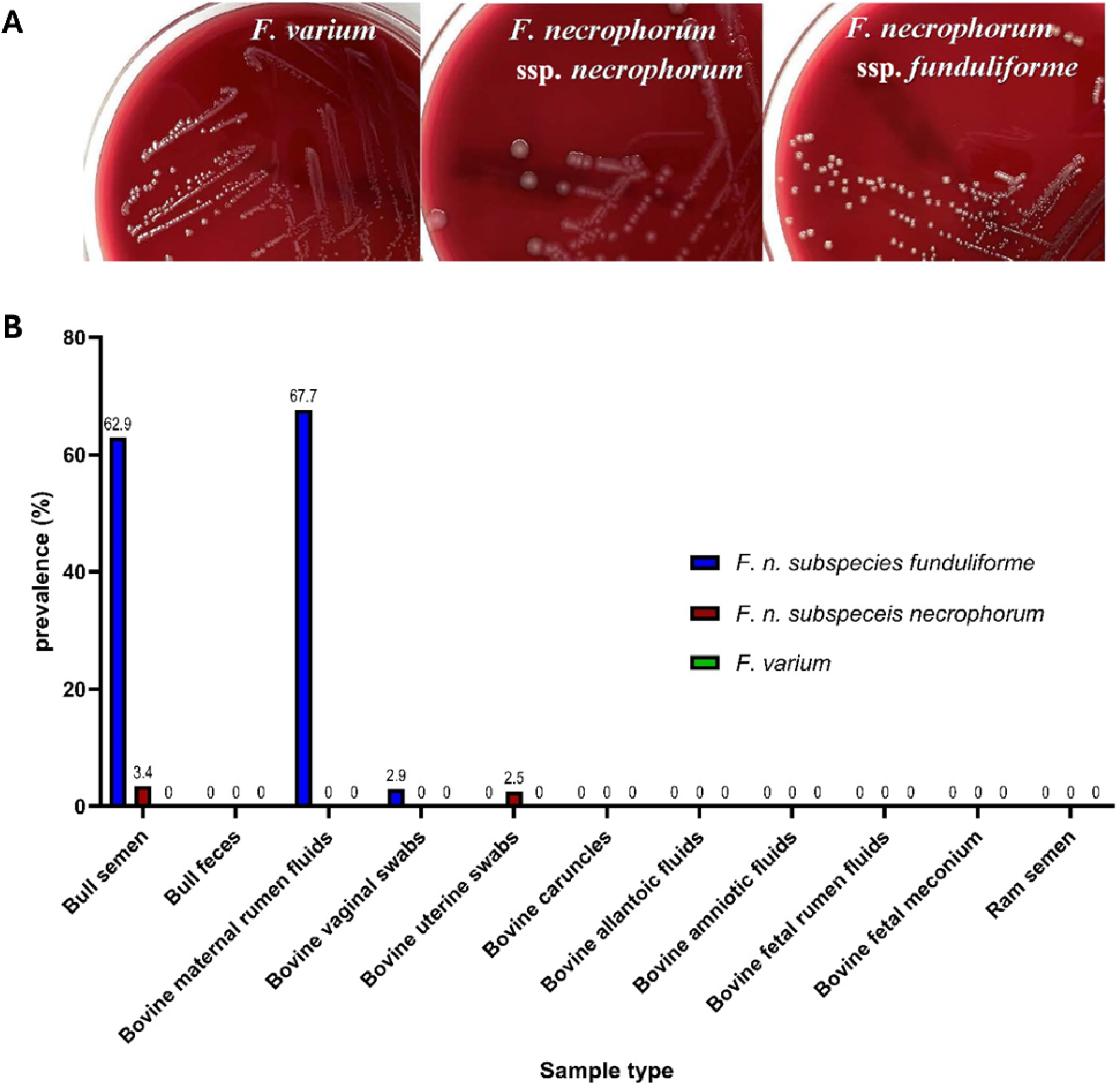
Targeted culturing of *Fusobacterium* spp. **(A)** The colony characteristics of *Fusobacterium varium, Fusobacterium necrophorum ssp. necrophorum,* and *ssp. funduliforme* isolated on blood agar media; Adopted from [22]; (**B**) Prevalence of targeted *Fusobacterium* spp. in different sample types (n = 499) as screened using targeted culturing techniques.

**Figure 4:**
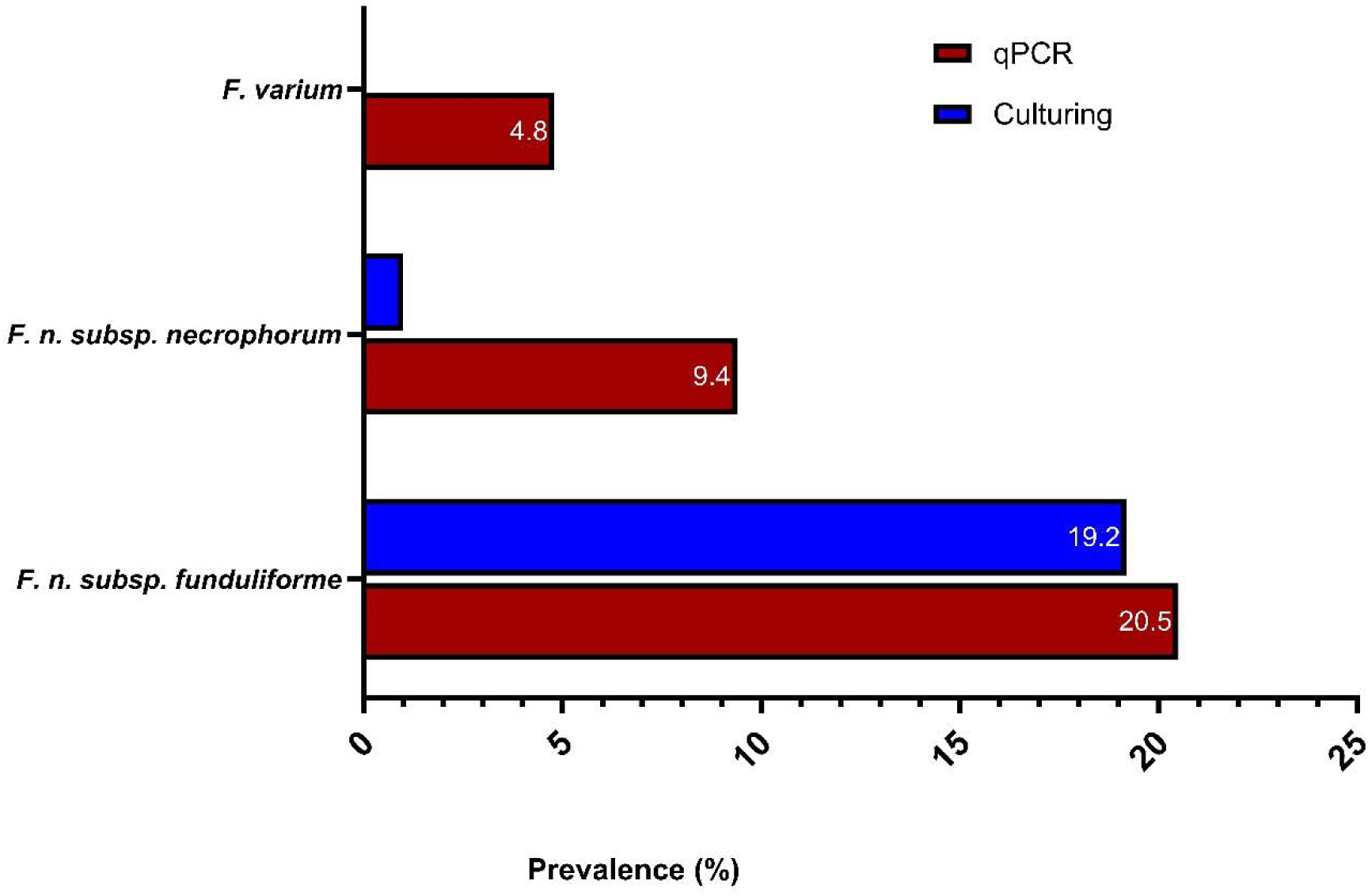
Overall prevalence of targeted *Fusobacterium* spp. in different sample types screened using both qPCR (n = 541) and targeted culturing (n = 499).

**Table 3:**
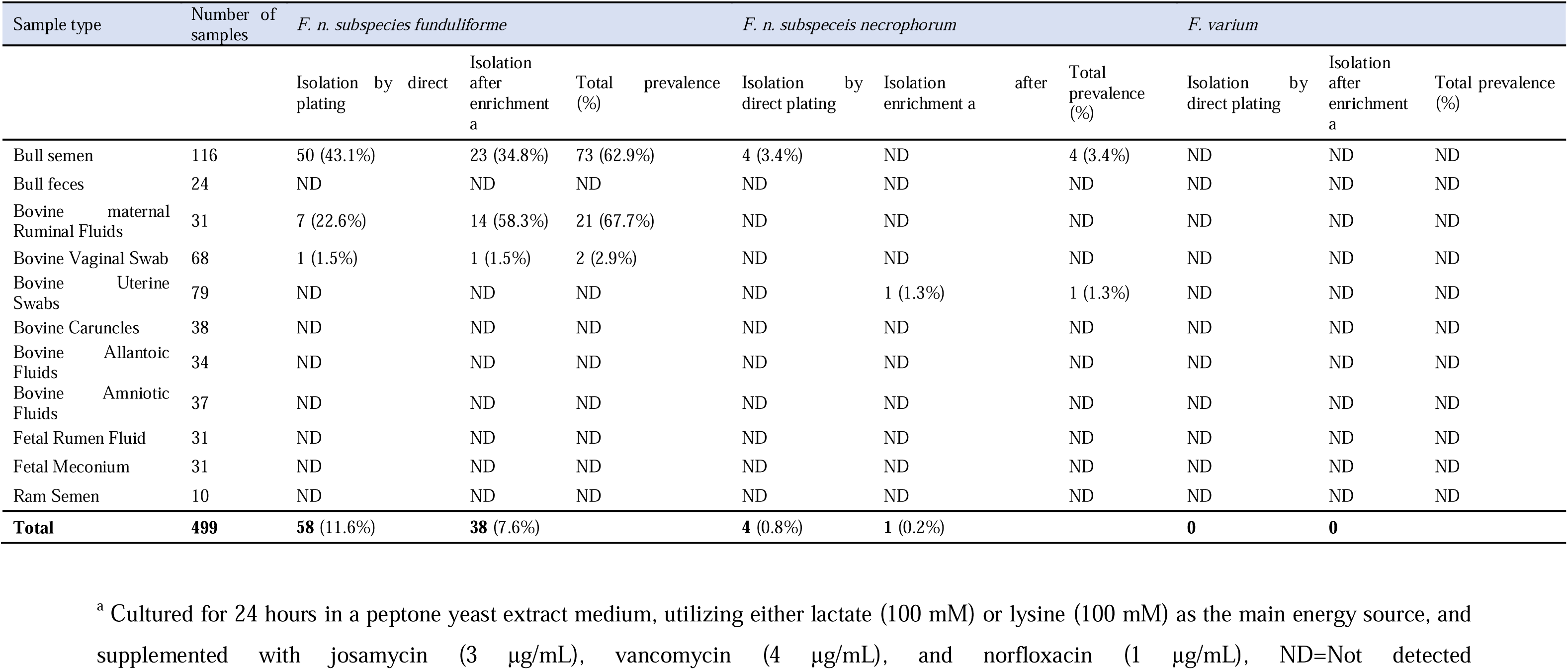
Frequency of isolation of targeted *Fusobacterium* spp. in different sample types both by direct plating and after enrichment.

Only four isolates of FNN were obtained from bull semen through direct plating, with no isolates recovered from post-enrichment cultures. Interestingly, a single FNN isolate was obtained from uterine swabs, but only after enrichment. No FNN was detected in all other sample types. FV was not isolated from any of the 499 samples analysed, neither by direct plating nor after enrichment.

## DISCUSSION

*Fusobacterium* species, particularly *F. necrophorum* and *F. varium,* colonize the gastrointestinal, respiratory, and reproductive tracts of both animals and humans where they are often associated with infections [10–13]. With its long reputation primarily as an opportunistic pathogen, *F necrophorum* is implicated in a variety of infections, such as liver abscesses, foot rot, laryngitis, and metritis in cattle [16, 43–46] as well as Lemierre’s syndrome in humans [47–49]. As opposed to its traditional view as a pathogen, our focus in the present study is to explore whether *Fusobacterium* subspecies and species are commensal members of the reproductive microbiota. We performed a comprehensive screening using both qPCR and targeted culturing to determine the prevalence of the *Fusobacterium* species; FNN, FNF, and FV in male (bovine and ovine) and female reproductive tracts, as well as other microbial ecosystems associated with the rumen and bovine calf fetuses.

Using the qPCR assay, we demonstrated that *Fusobacterium* spp. are variably prevalent in the semen of healthy bulls and rams with FNF being the most abundant subspecies. These findings align with previous studies where *Fusobacterium* was reported to be a dominant bacterial genus in the semen [28, 33, 34] and prepuce of healthy bulls [32], which was determined by 16S rRNA gene amplicon sequencing. Moreover, in our previous study using 16S rRNA gene amplicon sequencing, *Fusobacterium* was also found to be the most abundant genus in the seminal microbiota of healthy bulls, with a mean relative abundance of 26% [28]. Likewise, Medo and colleagues identified Fusobacteriota as the third most prevalent phylum, accounting for 18% of the phyla in the semen samples of Slovak Holstein Friesian breeding bulls [33]. Using a partitioning around medoids analysis, a clustering technique that uses actual data points, known as medoids, to select cluster centers, the authors determined that semen samples could be divided into two distinct clusters based on their microbiome compositions: one dominated by Actinobacteria and Firmicutes, and the other showing a high prevalence of Fusobacteriota [33]. Additionally, Koziol and colleagues reported that Fusobacteriota is one of the top bacterial phyla in the bull semen, along with Actinomycetota, Bacteroidota, Euryarchaeota, Bacillota, and Proteobacteria [28, 31]. This phylum was also observed in the microbiota associated with bull prepuce, and *Fusobacterium* is one of the relatively abundant genera in this community [32]. Furthermore, *Fusobacterium* has been reported to have strong associations with other key members of the bull semen associated bacterial genera such as *Bacteroides* and *Porphyromonas* [31]. Despite its relatively high abundance in the bull semen, previous studies have mainly focused on pathogenic effects of *Fusobacterium,* particularly in its association with sperm quality [31, 33, 50]. It is important to note that most of the available data on *Fusobacterium* spp. are derived from 16S rRNA gene amplicon sequencing, which cannot provide higher taxonomic resolution of *Fusobacterium* spp. at the species or subspecies levels —a limitation that our current culture-based study addressed. However, based on the results obtained from previous 16S rRNA gene amplicon sequencing studies, and our current qPCR and culturing results, it is reasonable to conclude the *Fusobacterium* spp., particularly FNF, are part of normal seminal microbial community, and may not merely be pathogenic. Thus, further research is warranted to study its role in male reproductive health and fertility.

Additionally, our research group previously observed a noticeable decline in the relative abundance of genus *Fusobacterium* in the seminal microbiota, with mean relative abundance dropping from 25.1% ± 1.3% (SE) prior to breeding to 18.04% ± 2.4% after 28 days of breeding season [28]. While the diet, age and environment (pasture vs. confinement housing) could contribute to the reduction in *Fusobacterium* abundance during breeding, the possible transfer of *Fusobacterium* from the bull semen to reproductive tract of heifers during mating could not be ruled out [35]. This finding is consistent with a previous human study on the complementary semino-vaginal microbiome in couples, where a notable reduction in the relative abundance of *Lactobacillus crispatus* was observed after copulation, accompanied by a strong similarity between the semen and vaginal microbiota [51]. Moreover, growing evidence supports the notion that seminal bacteria can “hitchhike” to the uterus, influencing the microbial landscape of the female reproductive tract [35]. Several studies in humans have demonstrated that microbes can traverse from the vagina through cervical mucous to the uterus [51–53]. A number of studies also emphasized the dynamic interplay between the seminal and vaginal microbiomes, proposing that the seminal bacteria like *Fusobacterium* spp. could integrate into the vaginal environment, creating a temporary but influential shift in microbial composition, which may influence uterine health, either contributing to dysbiosis or promoting a balanced microbiome [54, 55]. Accordingly, we proposed that the introduction of seminal microorganisms into the female reproductive system during mating could affect not only fertility and the development of the embryo, but also play a role in transferring paternal programming effects to the offspring [56].

In the present study, the qPCR analysis revealed the presence of *Fusobacterium* spp. in both the vaginal and uterine microbiota of healthy cattle, with a prevalence ranging from 3.1% to 9.8% across the samples screened. Importantly, we were also able to successfully isolate FNN from uterine swabs of healthy cows, providing direct evidence of its active colonization within the uterine environment. Interestingly, these results are consistent with our earlier study based on the 16S rRNA sequencing where *Fusobacterium* was a relatively abundant genus in the uterus of healthy beef heifers [29]. In addition, we observed that *F. necrophorum* was significantly more abundant in the uterine microbiota of cows that successfully conceived than those that remained open after artificial insemination [30]. Several 16S rRNA gene amplicon sequencing-based studies also reported that *Fusobacterium* is a consistent member of the healthy female reproductive microbiome in cattle although they did not provide deeper taxonomic identification [29, 30, 57]. *Fusobacterium* was detected in the vaginal microbiota of Brangus heifers, though no significant differences were observed between pregnant and non-pregnant groups [58]. Similarly, *Fusobacterium* was identified as a key genus in both the vaginal and cervical microbiota of healthy dairy heifers and cows, further suggesting *Fusobacterium* spp. as a beneficial member of the reproductive tract microbiota [59–61].

In the present study, FNF had the highest abundance among *Fusobacterium* spp. tested, likely due to its lower virulence compared to FNN, which is known to produce higher levels of leukotoxin [42, 62]. Moreover, it was reported that *Lactobacillus crispatus* demonstrate niche-specific adaptations, with unique genomic signatures enabling it to flourish in varying host environments such as humans and poultry [63]. These traits include variations in carbohydrate utilization, CRISPR-Cas immune systems, and prophage elements, reflecting the specific ecological shows clear niche-specific adaptations, with unique genomic signatures enabling it to flourish in varying host environments such as humans and poultry [63]. These traits include variations in carbohydrate utilization, CRISPR-Cas immune systems, and prophage elements, reflecting the specific ecological challenges of each host [63]. In a similar way, *Fusobacterium* spp. are likely to adapt to different host and body site niches, developing specialized genomic features that allow them to persist in environments like the bovine reproductive tract and fetal tissues. A comparative genomic study of different *F. necrophorum* strains revealed significant diversity in virulence genes, such as toxins, adhesion proteins, outer membrane proteins, cell envelope components, type IV secretion systems, ABC (ATP-binding cassette) transporters, and other transporter proteins, in addition to other distinct genomic signatures found among subspecies [13]. These differences highlight their adaptation to various environmental conditions and make it plausible to speculate that F. necrophorum strains in the reproductive tract may lack the virulence factors typically associated with pathogenic strains that cause conditions like liver abscesses, foot rot, endometritis, or abortion. The low expression of virulence could allow *Fusobacterium* spp. to be tolerated by immune cells within uterine ecological niches and potentially contribute to immunomodulation during pregnancy. Future studies should compare genomic profiles between *Fusobacterium* strains present in bovine reproductive tracts and those associated with infections and investigate the functional roles of virulence factors in different *Fusobacterium* spp. particularly in the context of fetal immune programming *in utero*.

Moreover, our recent study revealed that at the time of artificial insemination, the relative abundance of *Fusobacterium* in cows that later became pregnant was 125 times greater than in those that did not conceive [30] [30]. This body of evidence points to the possibility that *F. necrophorum,* particularly FNF, in the bovine urogenital tract could be more than just a pathogen; it may serve as a beneficial commensal with a positive role in promoting reproductive health and fertility. Moreover, *Fusobacteriota* in the maternal prenatal gut microbiota has been reported to be associated with enhanced fine motor skills in human fetus in gut microbiota of infant its associated with reduced fine motor skills, implying that these bacteria may have dual and opposing roles during different stages of fetal neurodevelopment [64].

However, the relatively low detection rate of *Fusobacterium* spp. in vaginal and uterine samples—ranging from 3.1% to 9.8% by qPCR and only 2.5% by targeted culture method still could suggest that *Fusobacterium* spp. is a not necessarily a consistent commensal in the bovine female urogenital microbiome. It is possible that *Fusobacterium* populations in the bovine female reproductive tract are transient, varying significantly depending on the reproductive cycle stage or immune status, which could lead to fluctuations in its presence and abundance in response to immunity [54, 65]. Another possible explanation could be that *Fusobacterium* spp. generally exists in low abundance in uterus [66] or perhaps present in a viable but non-culturable (VBNC) state within the reproductive tract [67, 68]. In the VBNC state, bacteria remain alive but do not grow on standard culture media, which might explain the low success rate of culturing even with enrichment steps, making them difficult to detect through culture methods. Another potential explanation for the low detection rates of *Fusobacterium* spp. in vaginal and uterine samples is that these bacteria, originating from the bovine female urogenital tract, may require specialized nutrients, growth factors (such as reproductive hormones), and optimized culturing conditions [69–71]. To address these limitations, the authors propose using shotgun metagenomic sequencing, a more sensitive next-generation technique, to characterize the vagino-uterine microbiota in longitudinal studies encompassing all pregnancy stages and dietary management. This approach could help determine whether *Fusobacterium* presence fluctuates with reproductive stages, physiological changes, or dietary management, offering deeper insights into its role as a beneficial commensal, a marker of reproductive health, or a potential contributor to fertility outcomes.

In the present study, we also identified a significant prevalence of FNN in various fetus-associated samples, including caruncles (maternal portion of the placenta) (26.3%), fetal ruminal fluids (9.7%), allantoic fluids (8.8%), maternal ruminal fluids and fetal meconium (both 6.5%), and amniotic fluids (2.7%). These findings are consistent with our earlier observations suggests that a diverse microbial community exists within the fetal environment, and they support the proposal that maternal factors, particularly nutrition, could play a crucial role in shaping the fetal microbiome [72]. This study also contributes to the growing body of evidence supporting the concept of *in utero* microbial colonization, a topic of increasing interest in reproductive biology and microbiology [5, 72–74]. Thus, the idea that the fetal environment is not sterile is continuously gaining a considerable traction and challenging the long-standing view of the womb as a microbe-free zone [72, 74]. Our current detection of FNN in both maternal and fetal compartments suggest a possible intra-uterine bacterial transfer from the mother to the fetus during gestation. Further research is needed to further verify the existence of prenatal colonization of *Fusobacterium* spp. and identify the routes and mechanisms by which these microbes are transmitted to the fetus, as well as their specific roles in reproductive health and fetal development. Additionally, this finding also implies that FNN, along with FNF and FV, may be integral to the microbial community that colonizes the fetus *in utero* that contribute to early microbial programming which might also affect the establishment and composition of the neonatal microbiome [75]. Notably, the presence of FNN across a range of fetus-associated samples, particularly in caruncles and fetal ruminal fluids, highlights its potential involvement in crucial physiological and immunological processes during gestation.

A recent study in humans examined the role of the prenatal gut microbiome in offspring neurodevelopment and reported that seven *Fusobacterium* amplicon sequence variants were shared between maternal and infant gut microbiomes, comprising 35% and 95% of their relative abundance, respectively, suggesting that vertical transmission influences their neurodevelopmental functions [64]. Several studies have proposed that early exposure to microbes play a role in developmental programming, possibly resulting in long-term impacts on the neonatal and infant gut microbiota and affecting overall health including metabolism, immunity, and disease resistance [40, 72, 76, 77]. In this context, the presence of FNN, FNF and FV in the bovine fetal environment could influence the development of the fetal immune system [77], potentially “educating” it in a manner that prepares the neonate for postnatal exposures *Fusobacterium* pathogens [78]. Additionally, it has recently been reported that bacterial extracellular vesicles in amniotic fluid resemble those from the maternal gut microbiota and potentially assist in priming the fetal immune system for postnatal gut colonization [78]. These were further supported by a recent human study which detected viable bacterial strains (e.g. *Staphylococcus* and *Lactobacillus* spp.) in fetal tissues such as gut, skin, lung, thymus, spleen, and placenta, which stimulated memory T cells in mesenteric lymph nodes of fetus *in vitro* [79]. This indicated that intrauterine microbial exposure may aid in priming the immunity and establishment of fetal immunocompetency before birth [79]. Many scholars have recently reviewed the vital role of early microbial interactions in shaping immune imprinting during both fetal development and postnatal life [56, 80–82]. They reported that compounds transferred from the maternal microbiota are crucial for the development of fetal innate immune cells, while the postnatal microbiota further supports the maturation of immune tissues and cells [80–82]. Thus, disruptions in interaction of maternal and fetal microbial processes could lead to pathological immune imprinting, increasing the risk of inflammatory conditions later in post-natal life [83]. Furthermore, a recent study revealed that human fetuses possess higher numbers of follicular and transitional B cells, along with CD4+ and CD8+ T cells with ability to secrete cytokines and display tissue-resident memory characteristics, which suggests that a complex innate and adaptive gut immune network develops before birth [84]. Therefore, this study, along with previous findings, underscores the possible role of microbial exposure in fetal development, suggesting that *Fusobacterium* species in the bovine uterus may play a role in shaping the fetal immune system and preparing it for postnatal microbial challenges. The potential involvement of *F. necrophorum* in fetal programming could shape the fetal immune development *in utero*, potentially determining their future susceptibility or resistance to infections like liver abscesses, mastitis, endometritis, and foot rot. Therefore, understanding possible involvement of *Fusobacterium* in immune priming is crucial for developing interventions that improve cattle health and resilience. Further research is warranted to investigate potential involvement of *F*. *necrophorum* in fetal immune development and later resistance against necrobacillosis, with the aim of refining management practices to effectively enhance cattle health and resilience.

Despite the high detection rates of *Fusobacterium* across various samples using qPCR, only 53.7% (101/188) of these detections were successfully isolated through both direct plating and enrichment, with the majority being FNF from bull semen and bovine ruminal fluids. This discrepancy could be stemming from the possibility that many *Fusobacterium* spp., particularly those detected in fetus-associated samples may be below the detection limit (< 10^3^ per ml or g) or in non-viable form, or they require special growth media and culturing conditions. Although they may not be viable in fetal environment, they still play a structural role in facilitating maternal-fetal microbiota interactions, which are believed to be crucial for priming the fetal immune system for postnatal gut colonization through bacterial vesicles [78]. Another factor potentially contributing to the discrepancy between qPCR detection and culturing results could be reduction in *Fusobacterium* viability because of exposure to aerobic conditions during sampling [85]. Fusobacteriota are obligatory anaerobes, meaning they thrive in environments devoid of oxygen, and exposure to aerobic conditions can significantly diminish their viability [86]. To mitigate this, samples were stored in pre-reduced 10% skim milk infused with carbon dioxide, with strict precautions taken to minimize oxygen exposure [87]. Skim milk is regarded as an effective cryopreservation medium, aiding in the preservation of bacterial stocks thawed from −80°C [88]. While efforts were made to plate the samples as fresh as possible, the geographical distance between the sampling sites and the microbiology laboratory necessitated freezing prior to plating. Hence, freezing the samples, which was a necessary step in this study due to logistical constraints, may have further impacted viability of some *Fusobacterium* spp. in the screened samples. Freezing is known to some extent adversely affect the viability and culturability of bacteria, and *Fusobacterium* spp. may exhibit varying degrees of tolerance to freezing and the cryoprotectants used [89]. Consequently, the culturing results may underrepresent the viable *Fusobacterium* species, particularly those susceptible to freeze-thaw cycles and oxygen exposure.

The limitations of this study, particularly in culturing *Fusobacterium* spp. due to potential aerobic exposure during sampling and freezing for transportation, were partially mitigated by adding an enrichment step to enhance bacterial recovery in the samples. While acknowledging the recent advancements and focus on metagenomic approaches [36, 90, 91], it is important to recognize that combination of both culture-dependent methods and molecular techniques continue to be indispensable for isolating and identifying bacteria at more precise taxonomic levels. These traditional methods provide essential insights into the morphological, genetic, and metabolic characteristics of bacteria, despite their time-consuming nature and the need for specific incubation conditions such as media composition, temperature, and oxygen levels [90]. Notably, the successful culturing of *Fusobacterium* spp. requires experience as well as specialized sample storage media and enrichment steps [21, 22]. It is important to highlight the robustness and comprehensiveness of this study, which integrated a diverse range of sample types from five distinct trials and employed both qPCR and culture-based methodologies to specifically target *Fusobacterium*. This approach contrasts with earlier studies that relied mainly on 16S rRNA sequencing which did not provide species and subspecies level resolution of *Fusobacterium* in the reproductive tract. In summary, among the *Fusobacterium* spp., FNF was the most prevalent subspecies, particularly in bull semen and bovine maternal ruminal fluids. Although enrichment techniques generally improved the isolation rates for FNF, they did not significantly enhance the recovery of FNN or enable the isolation of FV. These findings underscore the variability in the prevalence and culturing success across different *Fusobacterium* species and sample types, suggesting that alternative or optimized culturing conditions may be necessary to successfully recover FV and potentially other challenging species.

## CONCLUSIONS

We detected a high prevalence of FNF and FNN but relatively low prevalence of FV across various male and female reproductive tracts samples, including semen, vaginal, and uterine samples suggesting that these species may play beneficial roles in reproductive health and fertility. In addition, we also identified these *Fusobacterium* species in fetus-associated samples implying a possible transfer route from bull semen to cows, and subsequently to the developing fetus, potentially contributing to “priming” or “educating” the fetal immune system against future *Fusobacterium* associated infections. Overall, our findings call for further research to evaluate the potential role that *F.necrophorum* and other Fusobacterium spp. may have in cattle fertility and *in utero* fetal immune programming of calves.

## Acknowledgment

This study was funded by the North Dakota Agricultural Experiment Station as part of a start-up package for S.A. Mention of a trade name, proprietary product, or specific agreement does not constitute a guarantee or warranty by the USDA and does not imply approval to the inclusion of other products that may be suitable. USDA is an equal opportunity provider and employer.

## Author contributions

JK and SA conceptualized the study, designed the sampling strategy, and drafted the initial manuscript. SA, JK, CD, MC, RC, AS, KM, JC, and SA were responsible for sample collection. MA, XS, and TG handled sample processing and analyses. All authors contributed to the manuscript writing, revision, editing, and finalization, and approved the submitted version.

## Data availability

The data underlying this article will be shared on reasonable request to the corresponding author.

## REFERENCES

[1] L. P. Haraoui and M. J. Blaser, “The Microbiome and Infectious Diseases,” (in eng), Clin Infect Dis, vol. 77, no. Suppl 6, pp. S441–S446, Dec 05 2023, doi: 10.1093/cid/ciad577.

[2] E. Rackaityte and S. V. Lynch, “The human microbiome in the 21,” (in eng), Nat Commun, vol. 11, no. 1, p. 5256, Oct 16 2020, doi: 10.1038/s41467-020-18983-8.

[3] J. S. Leiby et al., “Lack of detection of a human placenta microbiome in samples from preterm and term deliveries,” (in eng), Microbiome, vol. 6, no. 1, p. 196, Oct 30 2018, doi: 10.1186/s40168-018-0575-4.

[4] S. Schoenmakers, R. Steegers-Theunissen, and M. Faas, “The matter of the reproductive microbiome,” (in eng), Obstet Med, vol. 12, no. 3, pp. 107–115, Sep 2019, doi: 10.1177/1753495X18775899.

[5] L. F. Stinson, M. C. Boyce, M. S. Payne, and J. A. Keelan, “The Not-so-Sterile Womb: Evidence That the Human Fetus Is Exposed to Bacteria Prior to Birth,” (in eng), Front Microbiol, vol. 10, p. 1124, 2019, doi: 10.3389/fmicb.2019.01124.

[6] N. K. Lema, M. T. Gemeda, and A. A. Woldesemayat, “Recent Advances in Metagenomic Approaches, Applications, and Challenge,” (in eng), Curr Microbiol, vol. 80, no. 11, p. 347, Sep 21 2023, doi: 10.1007/s00284-023-03451-5.

[7] U. Çömlekcioğlu, S. Jezierska, G. Opsomer, and O. B. Pascottini, “Uterine microbial ecology and disease in cattle: A review,” (in eng), Theriogenology, vol. 213, pp. 66–78, Jan 01 2024, doi: 10.1016/j.theriogenology.2023.09.016.

[8] S. Giannattasio-Ferraz et al., “A common vaginal microbiota composition among breeds of Bos taurus indicus (Gyr and Nellore),” (in eng), Braz J Microbiol, vol. 50, no. 4, pp. 1115–1124, Oct 2019, doi: 10.1007/s42770-019-00120-3.

[9] T. B. Ault et al., “Uterine and vaginal bacterial community diversity prior to artificial insemination between pregnant and nonpregnant postpartum cows1,” (in eng), J Anim Sci, vol. 97, no. 10, pp. 4298–4304, Oct 03 2019, doi: 10.1093/jas/skz210.

[10] T. G. Nagaraja, S. K. Narayanan, G. C. Stewart, and M. M. Chengappa, “Fusobacterium necrophorum infections in animals: pathogenesis and pathogenic mechanisms,” (in eng), Anaerobe, vol. 11, no. 4, pp. 239–46, Aug 2005, doi: 10.1016/j.anaerobe.2005.01.007.

[11] B. F. Langworth, “Fusobacterium necrophorum: its characteristics and role as an animal pathogen,” (in eng), Bacteriol Rev, vol. 41, no. 2, pp. 373–90, Jun 1977, doi: 10.1128/br.41.2.373-390.1977.

[12] T. G. Nagaraja and M. M. Chengappa, “Liver abscesses in feedlot cattle: a review,” (in eng), J Anim Sci, vol. 76, no. 1, pp. 287–98, Jan 1998, doi: 10.2527/1998.761287x.

[13] P. K. Bista, D. Pillai, C. Roy, J. Scaria, and S. K. Narayanan, “Comparative Genomic Analysis of Fusobacterium necrophorum Provides Insights into Conserved Virulence Genes,” (in eng), Microbiol Spectr, vol. 10, no. 6, p. e0029722, Dec 21 2022, doi: 10.1128/spectrum.00297-22.

[14] T. Shinjo, T. Fujisawa, and T. Mitsuoka, “Proposal of two subspecies of Fusobacterium necrophorum (Flügge) Moore and Holdeman: Fusobacterium necrophorum subsp. necrophorum subsp. nov., nom. rev. (ex Flügge 1886), and Fusobacterium necrophorum subsp. funduliforme subsp. nov., nom. rev. (ex Hallé 1898),” (in eng), Int J Syst Bacteriol, vol. 41, no. 3, pp. 395–7, Jul 1991, doi: 10.1099/00207713-41-3-395.

[15] S. Tadepalli, S. K. Narayanan, G. C. Stewart, M. M. Chengappa, and T. G. Nagaraja, “Fusobacterium necrophorum: a ruminal bacterium that invades liver to cause abscesses in cattle,” (in eng), Anaerobe, vol. 15, no. 1-2, pp. 36–43, 2009, doi: 10.1016/j.anaerobe.2008.05.005.

[16] R. G. Amachawadi and T. G. Nagaraja, “Pathogenesis of Liver Abscesses in Cattle,” (in eng), Vet Clin North Am Food Anim Pract, vol. 38, no. 3, pp. 335–346, Nov 2022, doi: 10.1016/j.cvfa.2022.08.001.

[17] I. Olsen, The Family Fusobacteriaceae. Springer, Berlin, Heidelberg, 2014.

[18] J. M. Donahue, Nonsporeforming Anaerobic Bacteria, Diagnostic Procedures in Veterinary Bacteriology and Mycology, Fifth Edition ed. Academic Press Inc., 1990.

[19] G. D. Bailey, Daria N., “Fusobacterium pseudonecrophorum Is a Synonym for Fusobacterium varium,” International Journal of Systematic and Evolutionary Microbiology, vol. 43, no. 4, pp. 819–821, 1993, doi: 10.1099/00207713-43-4-819.

[20] C. Schwarz et al., “Unexpected finding of Fusobacterium varium as the dominant Fusobacterium species in cattle rumen: potential implications for liver abscess etiology and interventions,” (in eng), J Anim Sci, vol. 101, Jan 03 2023, doi: 10.1093/jas/skad130.

[21] A. Deters, S. Xiaorong, B. Jianfa, K. Qing, M. Jacques, and T. G. Nagaraja, “A real-time PCR assay for the detection and quantification of Fusobacterium necrophorum and Fusobacterium varium in ruminal contents of cattle” Applied Animal Science, vol. 40, no. 3, pp. 250–259, 2024, doi: 10.15232/aas.2023-02507. Original Research Microorganisms and liver abscesses.

[22] A. Deters, X. Shi, L. Ty, and P. T. G. Nagaraja1, “First report of isolation of Fusobacterium varium from liver abscesses and ruminal and colonic epithelial tissues of feedlot cattle,” Applied Animal Science, vol. 40, no. 3, pp. 244–249, 2024, doi: 10.15232/aas.2023-02512. Original Research Microorganisms and liver abscesses.

[23] K. Clothier and M. Anderson, “Evaluation of bovine abortion cases and tissue suitability for identification of infectious agents in California diagnostic laboratory cases from 2007 to 2012,” (in eng), Theriogenology, vol. 85, no. 5, pp. 933–938, Mar 15 2016, doi: 10.1016/j.theriogenology.2015.11.001.

[24] I. M. Sheldon and H. Dobson, “Postpartum uterine health in cattle,” (in eng), Anim Reprod Sci, vol. 82-83, pp. 295–306, Jul 2004, doi: 10.1016/j.anireprosci.2004.04.006.

[25] D. E. Gomez, K. N. Galvão, J. C. Rodriguez-Lecompte, and M. C. Costa, “The Cattle Microbiota and the Immune System: An Evolving Field,” (in eng), Vet Clin North Am Food Anim Pract, vol. 35, no. 3, pp. 485–505, Nov 2019, doi: 10.1016/j.cvfa.2019.08.002.

[26] N. Mori et al., “Clinical characteristics and antimicrobial susceptibility of Fusobacterium species isolated over 10 years at a Japanese university hospital,” (in eng), Eur J Clin Microbiol Infect Dis, vol. 43, no. 3, pp. 423–433, Mar 2024, doi: 10.1007/s10096-023-04734-2.

[27] A. M. Almohaya, T. S. Almutairy, A. Alqahtani, K. Binkhamis, and F. M. Almajid, “Fusobacterium bloodstream infections: A literature review and hospital-based case series,” (in eng), Anaerobe, vol. 62, p. 102165, Apr 2020, doi: 10.1016/j.anaerobe.2020.102165.

[28] E. M. Webb et al., “A Longitudinal Characterization of the Seminal Microbiota and Antibiotic Resistance in Yearling Beef Bulls Subjected to Different Rates of Gain,” (in eng), Microbiol Spectr, vol. 11, no. 2, p. e0518022, Mar 14 2023, doi: 10.1128/spectrum.05180-22.

[29] T. M. Winders et al., “Feeding hempseed cake alters the bovine gut, respiratory and reproductive microbiota,” (in eng), Sci Rep, vol. 13, no. 1, p. 8121, May 19 2023, doi: 10.1038/s41598-023-35241-1.

[30] E. M. Webb et al., “Sequencing and culture-based characterization of the vaginal and uterine microbiota in beef cattle that became pregnant or remained open following artificial insemination,” (in eng), Microbiol Spectr, vol. 11, no. 6, p. e0273223, Dec 12 2023, doi: 10.1128/spectrum.02732-23.

[31] J. H. Koziol, T. Sheets, C. L. Wickware, and T. A. Johnson, “Composition and diversity of the seminal microbiota in bulls and its association with semen parameters,” (in eng), Theriogenology, vol. 182, pp. 17–25, Apr 01 2022, doi: 10.1016/j.theriogenology.2022.01.029.

[32] C. L. Wickware, T. A. Johnson, and J. H. Koziol, “Composition and diversity of the preputial microbiota in healthy bulls,” (in eng), Theriogenology, vol. 145, pp. 231–237, Mar 15 2020, doi: 10.1016/j.theriogenology.2019.11.002.

[33] J. Medo et al., “Core Microbiome of Slovak Holstein Friesian Breeding Bulls’ Semen,” (in eng), Animals (Basel), vol. 11, no. 11, Nov 22 2021, doi: 10.3390/ani11113331.

[34] A. Cojkic, A. Niazi, Y. Guo, T. Hallap, P. Padrik, and J. M. Morrell, “Identification of Bull Semen Microbiome by 16S Sequencing and Possible Relationships with Fertility,” (in eng), Microorganisms, vol. 9, no. 12, Nov 25 2021, doi: 10.3390/microorganisms9122431.

[35] S. M. Luecke, E. M. Webb, C. R. Dahlen, L. P. Reynolds, and S. Amat, “Seminal and vagino-uterine microbiome and their individual and interactive effects on cattle fertility,” (in eng), Front Microbiol, vol. 13, p. 1029128, 2022, doi: 10.3389/fmicb.2022.1029128.

[36] R. Koedooder et al., “Identification and evaluation of the microbiome in the female and male reproductive tracts,” (in eng), Hum Reprod Update, vol. 25, no. 3, pp. 298–325, May 01 2019, doi: 10.1093/humupd/dmy048.

[37] K. A. Bochantin-Winders et al., “Divergent planes of nutrition in mature rams influences body composition, hormone and metabolite concentrations, and offspring birth measurements, but not semen characteristics or offspring growth,” (in eng), J Anim Sci, Jul 24 2024, doi: 10.1093/jas/skae207.

[38] C. Kanjanaruch et al., “One-carbon metabolite supplementation to nutrient restricted beef heifers affects placental vascularity during early pregnancy,” vol. 102, ed. Oxford University Press, Great Clarendon Street, Oxford, OX2 6DP, United Kingdom: Journal of Animal Science, 2024, pp. 1–8.

[39] C. J. Kassetas et al., “Effects of feeding bulls dried corn distiller’s grains plus solubles or CaSO,” (in eng), Anim Reprod Sci, vol. 226, p. 106703, Mar 2021, doi: 10.1016/j.anireprosci.2021.106703.

[40] S. Amat et al., “Characterization of the Microbiota Associated With 12-Week-Old Bovine Fetuses Exposed to Divergent,” (in eng), Front Microbiol, vol. 12, p. 771832, 2021, doi: 10.3389/fmicb.2021.771832.

[41] M. S. Crouse et al., “Maternal nutrition and stage of early pregnancy in beef heifers: impacts on hexose and AA concentrations in maternal and fetal fluids1,” (in eng), J Anim Sci, vol. 97, no. 3, pp. 1296–1316, Mar 01 2019, doi: 10.1093/jas/skz013.

[42] F. Zhang, T. G. Nagaraja, D. George, and G. C. Stewart, “The two major subspecies of Fusobacterium necrophorum have distinct leukotoxin operon promoter regions,” (in eng), Vet Microbiol, vol. 112, no. 1, pp. 73–8, Jan 10 2006, doi: 10.1016/j.vetmic.2005.10.003.

[43] D. C. Van Metre, “Pathogenesis and Treatment of Bovine Foot Rot,” (in eng), Vet Clin North Am Food Anim Pract, vol. 33, no. 2, pp. 183–194, Jul 2017, doi: 10.1016/j.cvfa.2017.02.003.

[44] J. C. C. Silva et al., “Intrauterine infusion of a pathogenic bacterial cocktail is associated with the development of clinical metritis in postpartum multiparous Holstein cows,” (in eng), J Dairy Sci, vol. 106, no. 1, pp. 607–623, Jan 2023, doi: 10.3168/jds.2022-21954.

[45] D. K. Pillai, R. G. Amachawadi, G. Baca, S. Narayanan, and T. G. Nagaraja, “Leukotoxic activity of Fusobacterium necrophorum of cattle origin,” (in eng), Anaerobe, vol. 56, pp. 51–56, Apr 2019, doi: 10.1016/j.anaerobe.2019.02.009.

[46] D. K. Pillai, R. G. Amachawadi, G. Baca, S. K. Narayanan, and T. G. Nagaraja, “Leukotoxin production by Fusobacterium necrophorum strains in relation to severity of liver abscesses in cattle,” (in eng), Anaerobe, vol. 69, p. 102344, Jun 2021, doi: 10.1016/j.anaerobe.2021.102344.

[47] T. Riordan, “Human infection with Fusobacterium necrophorum (Necrobacillosis), with a focus on Lemierre’s syndrome,” (in eng), Clin Microbiol Rev, vol. 20, no. 4, pp. 622–59, Oct 2007, doi: 10.1128/CMR.00011-07.

[48] T. P. Atkinson et al., “Analysis of the tonsillar microbiome in young adults with sore throat reveals a high relative abundance of Fusobacterium necrophorum with low diversity,” (in eng), PLoS One, vol. 13, no. 1, p. e0189423, 2018, doi: 10.1371/journal.pone.0189423.

[49] Z. Mohiuddin, T. Manes, and A. Emerson, “Fusobacterium necrophorum Bacteremia With Evidence of Cavitary Pulmonary Lesion,” (in eng), Cureus, vol. 13, no. 11, p. e19537, Nov 2021, doi: 10.7759/cureus.19537.

[50] C. González-Marín et al., “Bacteria in bovine semen can increase sperm DNA fragmentation rates: a kinetic experimental approach,” (in eng), Anim Reprod Sci, vol. 123, no. 3-4, pp. 139–48, Feb 2011, doi: 10.1016/j.anireprosci.2010.11.014.

[51] R. Mändar et al., “Complementary seminovaginal microbiome in couples,” (in eng), Res Microbiol, vol. 166, no. 5, pp. 440–447, Jun 2015, doi: 10.1016/j.resmic.2015.03.009.

[52] L. K. Hansen et al., “The cervical mucus plug inhibits, but does not block, the passage of ascending bacteria from the vagina during pregnancy,” (in eng), Acta Obstet Gynecol Scand, vol. 93, no. 1, pp. 102–8, Jan 2014, doi: 10.1111/aogs.12296.

[53] R. Mändar, S. Türk, P. Korrovits, K. Ausmees, and M. Punab, “Impact of sexual debut on culturable human seminal microbiota,” (in eng), Andrology, vol. 6, no. 3, pp. 510–512, May 2018, doi: 10.1111/andr.12482.

[54] J. M. Baker, D. M. Chase, and M. M. Herbst-Kralovetz, “Uterine Microbiota: Residents, Tourists, or Invaders?,” (in eng), Front Immunol, vol. 9, p. 208, 2018, doi: 10.3389/fimmu.2018.00208.

[55] M. Benner, G. Ferwerda, I. Joosten, and R. G. van der Molen, “How uterine microbiota might be responsible for a receptive, fertile endometrium,” (in eng), Hum Reprod Update, vol. 24, no. 4, pp. 393–415, Jul 01 2018, doi: 10.1093/humupd/dmy012.

[56] J. Kilama, C. R. Dahlen, L. P. Reynolds, and S. Amat, “Contribution of the seminal microbiome to paternal programming,” (in eng), Biol Reprod, May 02 2024, doi: 10.1093/biolre/ioae068.

[57] C. T. Ong, C. Turni, P. J. Blackall, G. Boe-Hansen, B. J. Hayes, and A. E. Tabor, “Interrogating the bovine reproductive tract metagenomes using culture-independent approaches: a systematic review,” (in eng), Anim Microbiome, vol. 3, no. 1, p. 41, Jun 09 2021, doi: 10.1186/s42523-021-00106-3.

[58] R. D. Messman, Z. E. Contreras-Correa, H. A. Paz, G. Perry, and C. O. Lemley, “Vaginal bacterial community composition and concentrations of estradiol at the time of artificial insemination in Brangus heifers,” (in eng), J Anim Sci, vol. 98, no. 6, Jun 01 2020, doi: 10.1093/jas/skaa178.

[59] S. F. Lima, M. L. S. Bicalho, and R. C. Bicalho, “The Bos taurus maternal microbiome: Role in determining the progeny early-life upper respiratory tract microbiome and health,” (in eng), PLoS One, vol. 14, no. 3, p. e0208014, 2019, doi: 10.1371/journal.pone.0208014.

[60] J. J. Quereda et al., “Vaginal Microbiota Changes During Estrous Cycle in Dairy Heifers,” (in eng), Front Vet Sci, vol. 7, p. 371, 2020, doi: 10.3389/fvets.2020.00371.

[61] Y. Wang et al., “Characterization of the cervical bacterial community in dairy cows with metritis and during different physiological phases,” (in eng), Theriogenology, vol. 108, pp. 306–313, Mar 01 2018, doi: 10.1016/j.theriogenology.2017.12.028.

[62] G. Thapa, A. Jayal, E. Sikazwe, T. Perry, A. Mohammed Al Balushi, and P. Livingstone, “A genome-led study on the pathogenesis of Fusobacterium necrophorum infections,” (in eng), Gene, vol. 840, p. 146770, Oct 05 2022, doi: 10.1016/j.gene.2022.146770.

[63] M. Pan, C. Hidalgo-Cantabrana, and R. Barrangou, “Host and body site-specific adaptation of Lactobacillus Crispatus genomes,” (in eng), NAR Genom Bioinform, vol. 2, no. 1, p. lqaa001, Mar 2020, doi: 10.1093/nargab/lqaa001.

[64] Z. Sun et al., “Revealing the importance of prenatal gut microbiome in offspring neurodevelopment in humans,” (in eng), EBioMedicine, vol. 90, p. 104491, Apr 2023, doi: 10.1016/j.ebiom.2023.104491.

[65] C. G. Moreno, A. T. Luque, K. N. Galvão, and M. C. Otero, “Bacterial communities from vagina of dairy healthy heifers and cows with impaired reproductive performance,” (in eng), Res Vet Sci, vol. 142, pp. 15–23, Nov 19 2021, doi: 10.1016/j.rvsc.2021.11.007.

[66] L. Lietaer et al., “Low microbial biomass within the reproductive tract of mid-lactation dairy cows: A study approach,” (in eng), J Dairy Sci, vol. 104, no. 5, pp. 6159–6174, May 2021, doi: 10.3168/jds.2020-19554.

[67] J. Liu, L. Yang, B. V. Kjellerup, and Z. Xu, “Viable but nonculturable (VBNC) state, an underestimated and controversial microbial survival strategy,” (in eng), Trends Microbiol, vol. 31, no. 10, pp. 1013–1023, Oct 2023, doi: 10.1016/j.tim.2023.04.009.

[68] T. Ramamurthy, A. Ghosh, G. P. Pazhani, and S. Shinoda, “Current Perspectives on Viable but Non-Culturable (VBNC) Pathogenic Bacteria,” (in eng), Front Public Health, vol. 2, p. 103, 2014, doi: 10.3389/fpubh.2014.00103.

[69] E. Rackaityte et al., “Viable bacterial colonization is highly limited in the human intestine in utero,” (in eng), Nat Med, vol. 26, no. 4, pp. 599–607, Apr 2020, doi: 10.1038/s41591-020-0761-3.

[70] E. K. Mallott, C. Borries, A. Koenig, K. R. Amato, and A. Lu, “Reproductive hormones mediate changes in the gut microbiome during pregnancy and lactation in Phayre’s leaf monkeys,” (in eng), Sci Rep, vol. 10, no. 1, p. 9961, Jun 19 2020, doi: 10.1038/s41598-020-66865-2.

[71] A. Canha-Gouveia et al., “Physicochemical and Functional Characterization of Female Reproductive Fluids: A Report of the First Two Infants Born Following Addition of Their Mother’s Fluids to the Embryo Culture Media,” (in eng), Front Physiol, vol. 12, p. 710887, 2021, doi: 10.3389/fphys.2021.710887.

[72] S. Amat, C. R. Dahlen, K. C. Swanson, A. K. Ward, L. P. Reynolds, and J. S. Caton, “Bovine Animal Model for Studying the Maternal Microbiome,” (in eng), Front Microbiol, vol. 13, p. 854453, 2022, doi: 10.3389/fmicb.2022.854453.

[73] C. E. Guzman et al., “A pioneer calf foetus microbiome,” (in eng), Sci Rep, vol. 10, no. 1, p. 17712, Oct 19 2020, doi: 10.1038/s41598-020-74677-7.

[74] G. Hummel and K. Aagaard, “Arthropods to Eutherians: A Historical and Contemporary Comparison of Sparse Prenatal Microbial Communities Among Animalia Species,” (in eng), Am J Reprod Immunol, vol. 92, no. 2, p. e13897, Aug 2024, doi: 10.1111/aji.13897.

[75] M. S. Vidal and R. Menon, “In utero priming of fetal immune activation: Myths and mechanisms,” (in eng), J Reprod Immunol, vol. 157, p. 103922, Jun 2023, doi: 10.1016/j.jri.2023.103922.

[76] D. D. Nyangahu et al., “Disruption of maternal gut microbiota during gestation alters offspring microbiota and immunity,” (in eng), Microbiome, vol. 6, no. 1, p. 124, Jul 07 2018, doi: 10.1186/s40168-018-0511-7.

[77] D. D. Nyangahu and H. B. Jaspan, “Influence of maternal microbiota during pregnancy on infant immunity,” (in eng), Clin Exp Immunol, vol. 198, no. 1, pp. 47–56, Oct 2019, doi: 10.1111/cei.13331.

[78] A. Kaisanlahti et al., “Maternal microbiota communicates with the fetus through microbiota-derived extracellular vesicles,” (in eng), Microbiome, vol. 11, no. 1, p. 249, Nov 13 2023, doi: 10.1186/s40168-023-01694-9.

[79] A. Mishra et al., “Microbial exposure during early human development primes fetal immune cells,” (in eng), Cell, vol. 184, no. 13, pp. 3394–3409.e20, Jun 24 2021, doi: 10.1016/j.cell.2021.04.039.

[80] Z. Al Nabhani and G. Eberl, “Imprinting of the immune system by the microbiota early in life,” (in eng), Mucosal Immunol, vol. 13, no. 2, pp. 183–189, Mar 2020, doi: 10.1038/s41385-020-0257-y.

[81] K. Donald and B. B. Finlay, “Early-life interactions between the microbiota and immune system: impact on immune system development and atopic disease,” (in eng), Nat Rev Immunol, vol. 23, no. 11, pp. 735–748, Nov 2023, doi: 10.1038/s41577-023-00874-w.

[82] X. Zhang, D. Zhivaki, and R. Lo-Man, “Unique aspects of the perinatal immune system,” (in eng), Nat Rev Immunol, vol. 17, no. 8, pp. 495–507, Aug 2017, doi: 10.1038/nri.2017.54.

[83] P. Henneke, K. Kierdorf, L. J. Hall, M. Sperandio, and M. Hornef, “Perinatal development of innate immune topology,” (in eng), Elife, vol. 10, May 25 2021, doi: 10.7554/eLife.67793.

[84] S. F. Stras et al., “Maturation of the Human Intestinal Immune System Occurs Early in Fetal Development,” (in eng), Dev Cell, vol. 51, no. 3, pp. 357–373.e5, Nov 04 2019, doi: 10.1016/j.devcel.2019.09.008.

[85] S. Hoffmann and T. Justesen, “Effect of temperature, humidity and exposure to oxygen on the survival of anaerobic bacteria,” (in eng), J Med Microbiol, vol. 13, no. 4, pp. 609–12, Nov 1980, doi: 10.1099/00222615-13-4-609.

[86] Z. Lu and J. A. Imlay, “When anaerobes encounter oxygen: mechanisms of oxygen toxicity, tolerance and defence,” (in eng), Nat Rev Microbiol, vol. 19, no. 12, pp. 774–785, Dec 2021, doi: 10.1038/s41579-021-00583-y.

[87] S. Reilly, “The carbon dioxide requirements of anaerobic bacteria,” (in eng), J Med Microbiol, vol. 13, no. 4, pp. 573–9, Nov 1980, doi: 10.1099/00222615-13-4-573.

[88] W. L. Cody et al., “Skim milk enhances the preservation of thawed −80 degrees C bacterial stocks,” (in eng), J Microbiol Methods, vol. 75, no. 1, pp. 135–8, Sep 2008, doi: 10.1016/j.mimet.2008.05.006.

[89] L. Bircher, A. Geirnaert, F. Hammes, C. Lacroix, and C. Schwab, “Effect of cryopreservation and lyophilization on viability and growth of strict anaerobic human gut microbes,” (in eng), Microb Biotechnol, vol. 11, no. 4, pp. 721–733, Jul 2018, doi: 10.1111/1751-7915.13265.

[90] P. C. Woo, S. K. Lau, J. L. Teng, H. Tse, and K. Y. Yuen, “Then and now: use of 16S rDNA gene sequencing for bacterial identification and discovery of novel bacteria in clinical microbiology laboratories,” (in eng), Clin Microbiol Infect, vol. 14, no. 10, pp. 908–34, Oct 2008, doi: 10.1111/j.1469-0691.2008.02070.x.

[91] Z. Al-Kass, E. Eriksson, E. Bagge, M. Wallgren, and J. M. Morrell, “Microbiota of semen from stallions in Sweden identified by MALDI-TOF,” (in eng), Vet Anim Sci, vol. 10, p. 100143, Dec 2020, doi: 10.1016/j.vas.2020.100143.

